# Identification of biomarkers and pathways to identify novel therapeutic targets in Alzheimer’s disease: Insights from a systems biomedicine perspective

**DOI:** 10.1101/481879

**Authors:** Md. Rezanur Rahman, Tania Islam, Toyfiquz Zaman, Md. Shahjaman, Md. Rezaul Karim, Fazlul Huq, Julian M.W. Quinn, R. M. Damian Holsinger, Mohammad Ali Moni

## Abstract

Alzheimer’s disease (AD) is a progressive neurodegenerative disease characterized by the accumulation of amyloid plaques and neurofibrillary tangles in the brain. However, there are no peripheral biomarkers available that can detect AD onset. This study aimed to identify such AD biomarkers through an integrative analysis of blood cell gene expression data. We used two microarray datasets (GSE4226 and GSE4229) comparing blood cell transcriptomes of AD patients and matched controls to identify differentially expressed genes (DEGs). Geneset and protein overrepresentation analysis, protein-protein interaction (PPI), DEGs-Transcription Factor interactions, DEGs-MicroRNAs interactions, protein-drug interactions, and protein subcellular localizations analyses were performed on DEGs common to the datasets. We identified 25 common DEGs between the two datasets. Integration of genome scale transcriptome datasets with biomolecular networks revealed hub genes (TUBB, ATF3, NOL6, UQCRC1, SND1, CASP2, BTF3, INPP5K, VCAM1, and CSTF1), common transcription factors (FOXC1, ZNF3, GEMIN7, and SMG9) and miRNAs (mir-20a-5p, mir-93-5p, mir-16-5p, let-7b-5p, mir-708-5p, mir-24-3p, mir-26b-5p, mir-17-5p, mir-4270, and mir-4441). These included 8 previously identified AD markers (BTF3, VCAM, FOXC1, mir-26b-5p, mir-20a-5p, mir-93-5p, let-7b-5p, mir-24-3p) and 13 novel AD biomarkers. Evaluation of histone modification revealed that hub genes and transcription factors possess several histone modification sites associated with AD. Protein-drug interactions revealed 10 compounds that affect the identified AD candidate markers, including anti-neoplastic agents (Vinorelbine, Vincristine, Vinblastine, Epothilone D, Epothilone B, CYT997, and ZEN-012), a dermatological (Podofilox) and an immunosuppressive agent (Colchicine). The subcellular localization of markers varied, including nuclear, plasma membrane and cytosolic proteins. This study identified blood-cell derived molecular biomarker signatures that might be useful as peripheral biomarkers in AD. This study also identified drug and epigenetic data associated with these molecules that may be useful in designing therapeutic approaches to ameliorate AD.

## 1. Introduction

Alzheimer’s disease (AD) is a progressive neurodegenerative disease that gives rise to dementia and severe impairment of cognitive function in affected people, and is notably associated with the formation of extracellular amyloid plaques and intracellular accumulation of neurofibrillary tangles in the brain. It is the most common form of dementia, affecting 5.5 million individuals in the US alone, (Association A, 2017). Cognitive impairment is the main assessment method for patients, [2]. Sporadic AD is the most common form of the disease and generally affects individuals over the age of 65. Early diagnosis is rare, occurring in only a small fraction (1-5%) of patients. Early diagnosis of AD may improve management strategies for patients. However, there are no effective treatments for the disease. Therefore, research directed towards identifying AD biomarkers is needed not just for a better understanding of the development of AD but also as an important step in discovering new treatment strategies [3].

For differential diagnosis and AD patient assessments, neuroimaging techniques and cerebrospinal fluid biomarkers are widely used in clinical practice [4–6]. These methods are expensive, show low sensitivity and specificity and in the case cerebrospinal fluid, highly invasive. Thus, a simple blood-based test of dementia patients would be a valuable clinical tool for AD [7]. However, considering the dearth of useful peripheral blood protein or antigen biomarkers, despite much effort [8–11], blood-cell based transcript biomarkers may be a valuable approach [12–16]. Indeed, epigenetic factors are known to affect the pathogenesis of AD and have been accepted as playing significant roles in the development and progression of neurodegenerative diseases. Such epigenetic profiling, including studies of DNA methylation and histone modifications can be used to reveal the regulation of gene expression in various diseases [17]. Various external factors such as lifestyle, age, environment and health status result in epigenetic and transcriptome changes that may relate to AD development [17]. However, epigenetic studies have yet to identify key mechanisms in AD development [17].

Dysregulation of the miRNAs has also been implicated in AD [18] and, consequently, miRNAs are increasingly being examined as possible biomarkers in or mediating factors in AD [19–21]. Thus, miRNA and epigenetic approaches have led to the use of systems biology approaches to elucidate key mediating or marker biomolecules in different complex diseases such as AD [20–23]. Accordingly, we have employed such an approach here. To identify critical genes, we used two peripheral blood microarray gene expression datasets derived from studies of AD patients. Using DEGs identified by this approach we performed functional annotations to identify important Gene Ontology and pathways enriched in these DEGs. We integrated the DEGs with interaction networks using the following approaches:

(i) a PPI network of the proteins encoded by DEGs;
(ii) DEG-TFs and DEGs-miRNAs interactions;
(iii) protein-drug interaction networks to screen potential drugs with the binding affinity to target proteins estimated by molecular docking simulations;
(iv) prediction of subcellular localization of proteins encoded by the DEGs was performed with the intent to provide insights into function.

Thus, we have used a comprehensive systems biology pipeline to explore molecular biomarker signatures of both mRNAs and miRNAs present in peripheral blood transcriptomes of AD patients. Additionally, we have also identified potential hub proteins, transcription factors (TFs) and pathways in blood that may provide potential biomarkers for early diagnosis of AD. These discoveries provide novel insights into the mechanisms underlying AD (Figure 1).

**Figure 1:**
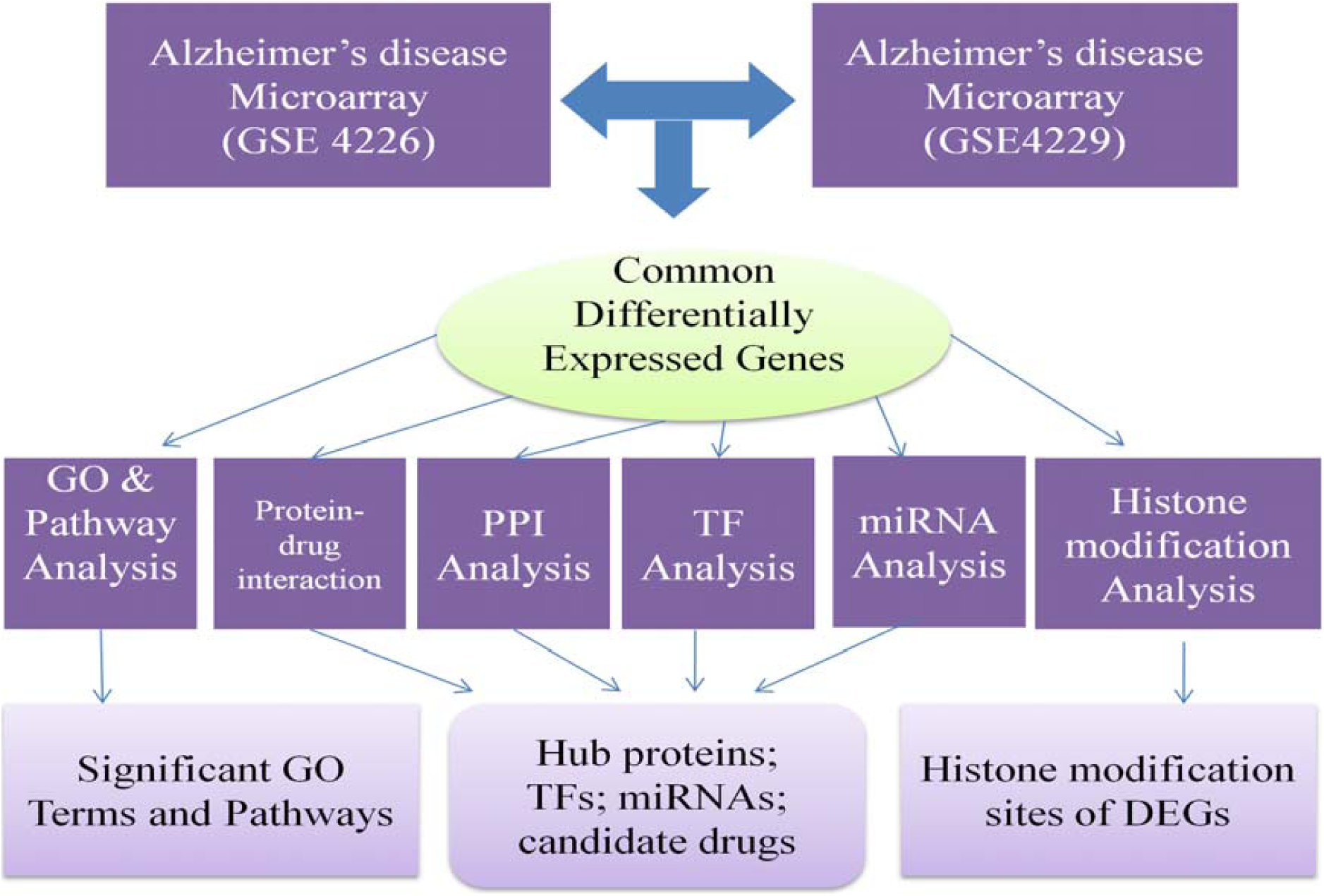
A multi-stage analysis methodology employed in this study. Transcriptomic datasets of blood from AD and normal individuals were collected from the NCBI-GEO database. The datasets were analyzed using GEO2R to identify DEGs. Functional enrichment analyses of DEGs were performed to identify significantly enriched pathways, Gene Ontology terms, and disease overrepresentation terms. Protein-protein interaction networks were reconstructed around DEGs. TF-target gene interactions and miRNA-target gene interactions were downloaded from JASPAR and miRTarbase databases, respectively. The protein-drug interaction analysis using the DrugBank database revealed candidate small molecular drugs that may interact with identified proteins. Significant pathways and GO terms, candidate biomarkers at protein and TFs levels, drug target, candidate drugs and drug-targeting sites were also identified.

## 2. Materials and Methods

### 2.1 Identification of Differentially Expressed Genes from Microarray High-throughput Datasets of Blood of AD Patients

The microarray gene expression high-throughput datasets in AD was obtained from the NCBI-GEO database [24]. This data was obtained from two separate studies of peripheral blood mononuclear cells that were collected from normal elderly control and AD subjects and analysed using human MGC cDNA microarrays; the results were deposited in NCBI-GEO database with the accession numbers GSE4226 and GSE4229 [25]. GSE4226 contained samples from 28 individuals in total, including 7 each of male and female controls and male and female AD patients, with the analysis originally published by Maes et al. [14] [PMID: 19366883], who also used DNA and blood protein studies in the search for new biomarkers. GSE4229 studied 40 blood samples (11 from male controls, 12 from female controls, 10 from male AD patients and 7 from female AD patients). The datasets were originally analyzed by Maes and colleagues using the NCBI-GEO2R online tool to identify DEGs in the AD compared to age matched controls. In our analysis, the gene expression dataset was first normalized by log2 transformation. Then, we analyzed the datasets in NCBI’s GEO2R tool using Limma in hypothesis testing and the Benjamini & Hochberg correction to control the false discovery rate. The p-value<0.01 was regarded as the cut-off criteria to screen for significant DEGs. Venn analysis was performed using the online tool–Jvenn [26] to identify common target DEGs from the two datasets.

### 2.2 Gene Ontology and Pathway Enrichment Analysis

Gene over-representation analyses were performed via the bioinformatics resource Enrichr [27] to identify molecular function, biological process, and pathway annotations of the identified DEGs. In these analyses, the Gene Ontology (GO) terminology and KEGG pathway database were used as annotation sources. The protein class over-representation was performed using the PANTHER database [28]. A p-value<0.05 was considered significant for all enrichment analyses.

### 2.3 Protein-protein Interaction Analysis

We used the STRING protein interactome database [29] to construct PPI networks of the proteins encoded by the DEGs. We set a medium confidence score of 400 for the construction of PPI networks since the number of DEGs is low. Network visualization and topological analyses were performed using NetworkAnalyst (Xia et al., 2015).

### 2.4 DEG-TFs Interaction Analysis

To identify regulatory TFs that control the DEGs at a transcriptional level, TF-target gene interactions were obtained using the JASPAR database [31] and were based on topological parameters (i.e., degree and betweenness centrality) using NetworkAnalyst [30].

### 2.5 DEGs-miRNA Interaction Analysis

To identify regulatory miRNAs that influence DEGs at the post-transcriptional level, miRNAs-target gene interactions were obtained from TarBase [32] and miRTarBase [33] based on topological parameters (i.e., degree and betweenness centrality) and analyzed using NetworkAnalyst [30].

### 2.6 Identification of Histone Modification Sites

Based on the recent systematic review on neurodegenerative disease aimed to identify most consistent epigenetic marks reported in methylations genes and histone modifications associated with AD [34], we used the human histone modification database (HHMD) to identify histone modification sites of the hub genes and TFs [35].

### 2.7 Cross-Validation of Candidate Biomarker Biomolecules

The AlzGene database contains 618 published genetic association studies of AD [36] and was used to evaluate the interactions between AD-associated cellular alterations in blood and AD-associated alterations collected in AlzGene database. The miRNAs were also crosschecked with blood-based miRNA signatures.

### 2.8 Protein-drug Interactions Analysis

Protein-drug interaction was analyzed using the DrugBank database (Version 5.0) [37] via NetworkAnalyst to identify potential drugs that could be used in the treatment of AD. The protein-drug interaction analysis revealed that TUBB had interactions with several drugs. Then, in order perform molecular docking analysis between TUBB and drugs, three dimensional (3D) crystal structure of the target proteins TUBB (PDB ID:1TUB) was obtained from protein data bank (PDB) [38]. Molecular docking analyses were performed using the protein-small molecule docking server SwissDock [39]. The docking score and best-fit pose were selected for each ligand/drug.

### 2.9 Prediction of Protein Subcellular Localization

We used the WoLF PSORT software package [40] to predict the subcellular localizations of the proteins encoded by the DEGs. The WoLF PSORT software predicts subcellular localizations based on amino acid sequence information.

## 3. Results

### 3.1 Transcriptomic Signatures of AD

The microarray datasets obtained from blood tissue of AD patients were analyzed and 25 mutually expressed core DEGs were identified between the two datasets. These core DEGs were considered as transcriptomic signatures for AD (Figure 2A) and classified into groups according to their functions and activities as transporters (4%), membrane traffic proteins (4%), hydrolases (24%), oxidoreductases (4%), cell junction proteins (4%), enzyme modulators (8%), transcription factors (12%), nucleic acid binding proteins (12%), calcium-biding proteins (4%) and cytoskeletal proteins (4%) (Figure 2B). A gene-set enrichment analysis showed that the DEGs were enriched in different biological processes, molecular functions and cellular components as summarized in Table 1. The molecular pathway enrichment analysis revealed that pathways previously identified in leukocyte transendothelial migration, oxidative phosphorylation, Parkinson’s disease, cell adhesion molecules, non-alcoholic fatty liver disease, Alzheimer’s disease and Huntington’s disease were altered (Table 2).

**Table 1:**
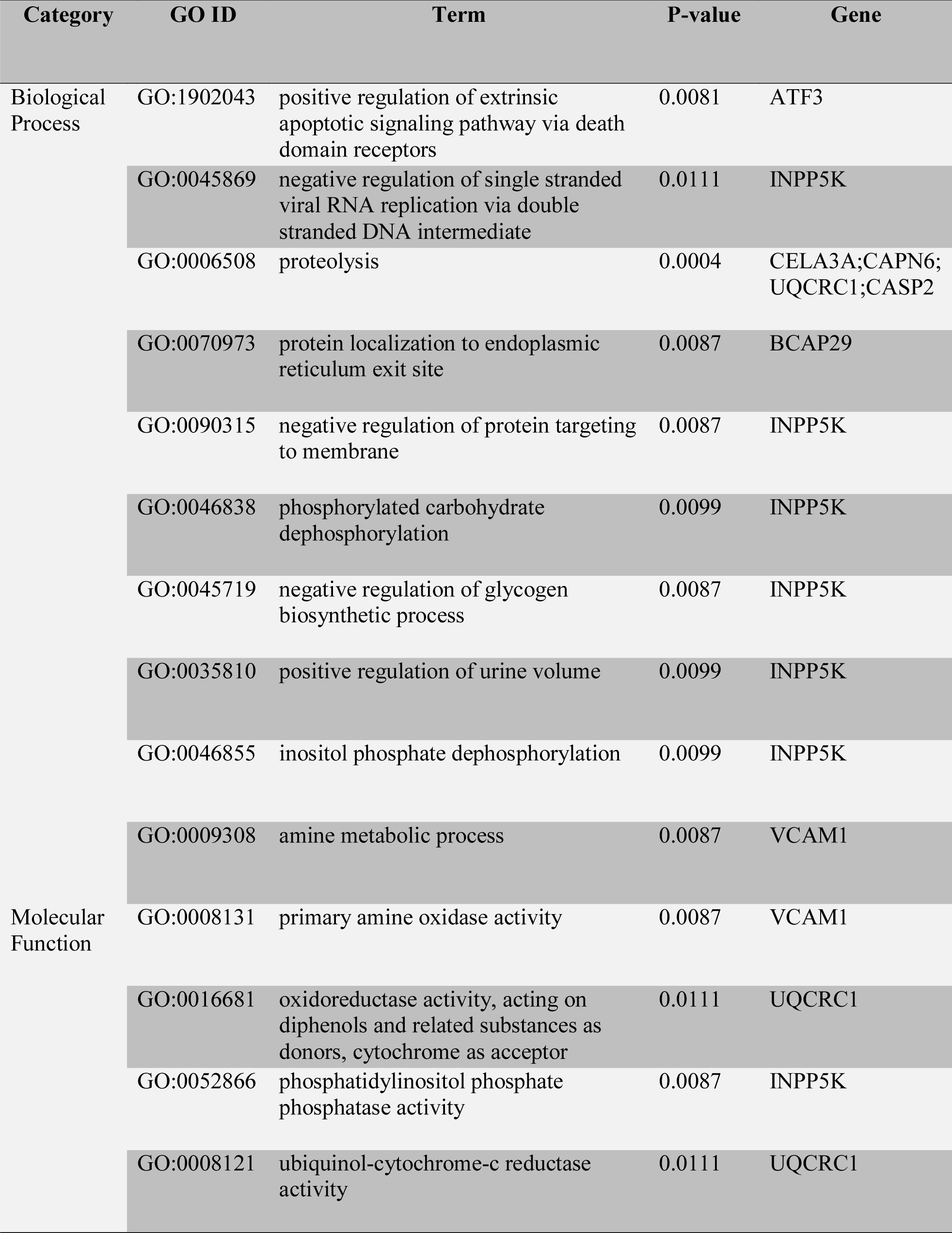
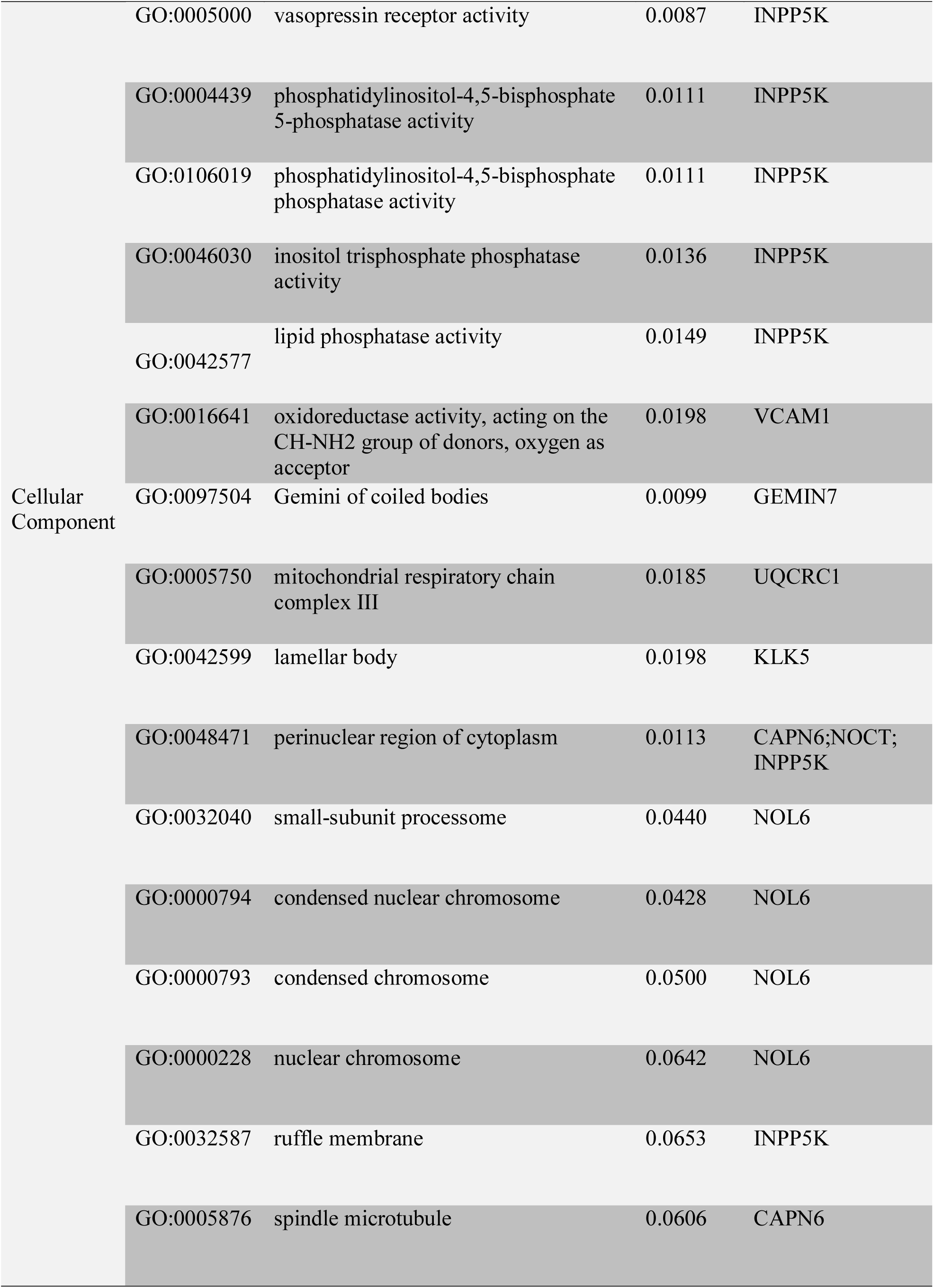
Gene set enrichment analysis of the differentially expressed genes identified from microarray data of blood of AD patients

**Table 2:** The molecular pathway enrichment in Alzheimer’s disease.

| Category | Pathways | Adj. p-value | Genes |
| --- | --- | --- | --- |
| KEGG | Ribosome | 0.021 | RPS9;RPL12;RPS5 |
| BioCarta | Alternative Complement Pathway | 0.02 | C3 |
|  | Classical Complement Pathway | 0.02 | C3 |
|  | Lectin Induced Complement Pathway | 0.02 | C3 |
| WikiPathways | Cytoplasmic Ribosomal Proteins | 0.005 | RPS9;RPL12;RPS5 |

**Figure 2:**
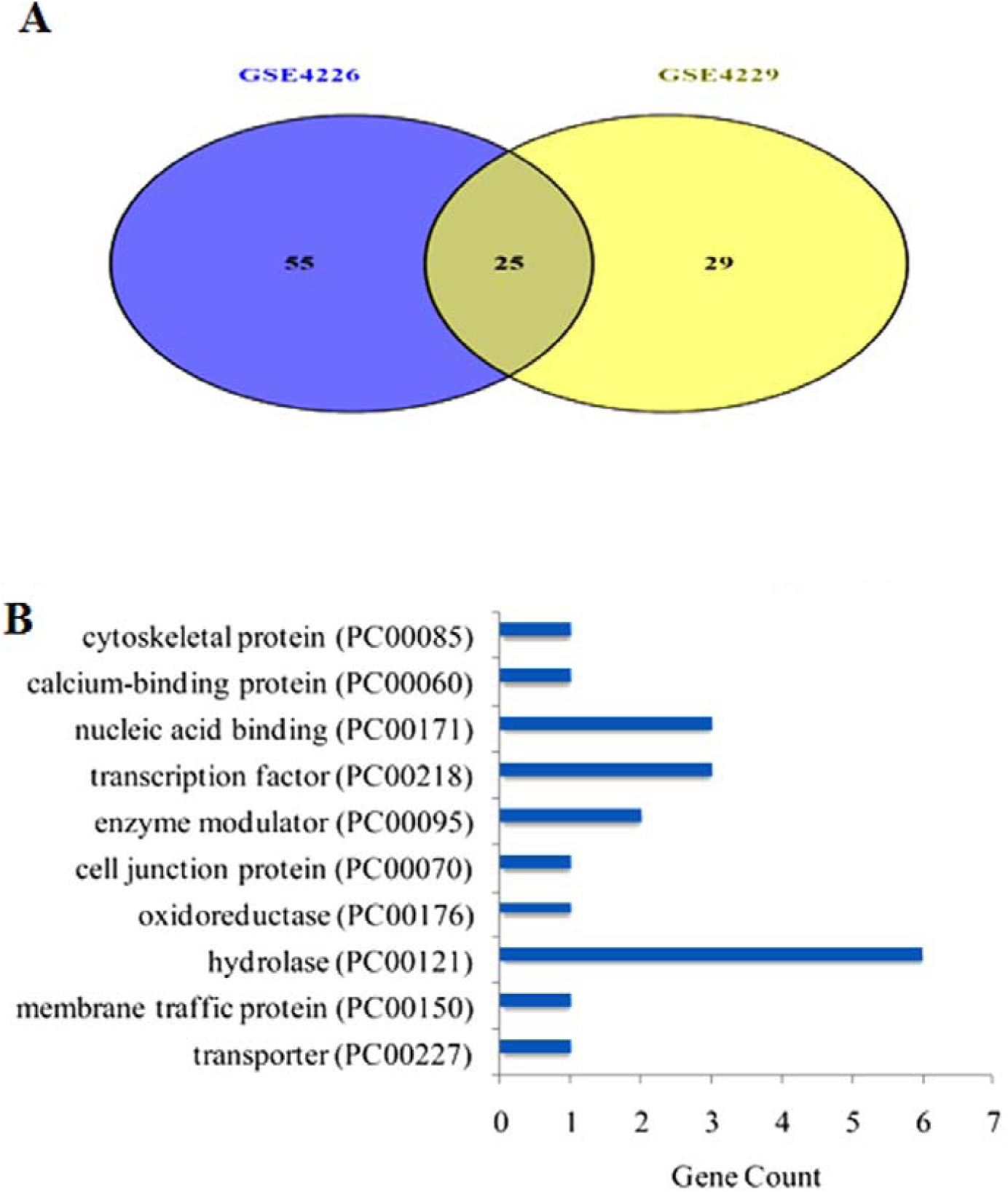
(A) Identification of DEGs in microarray datasets GSE4226 and GSE4229. (A) The mutually expressed core DEGs identified between two datasets. (B) The protein class over-representation coded by the DEGs identified using the PANTHER database.

### 3.3 Proteomic Signatures in AD

To reveal central proteins, a protein-protein interaction of the DEGs was constructed (Figure 3). Topological analysis revealed 10 central hub proteins (TUBB, ATF3, NOL6, UQCRC1, SND1, CASP2, BTF3, INPP5K, VCAM1, and CSTF1) (Table 3).

**Table 3:**
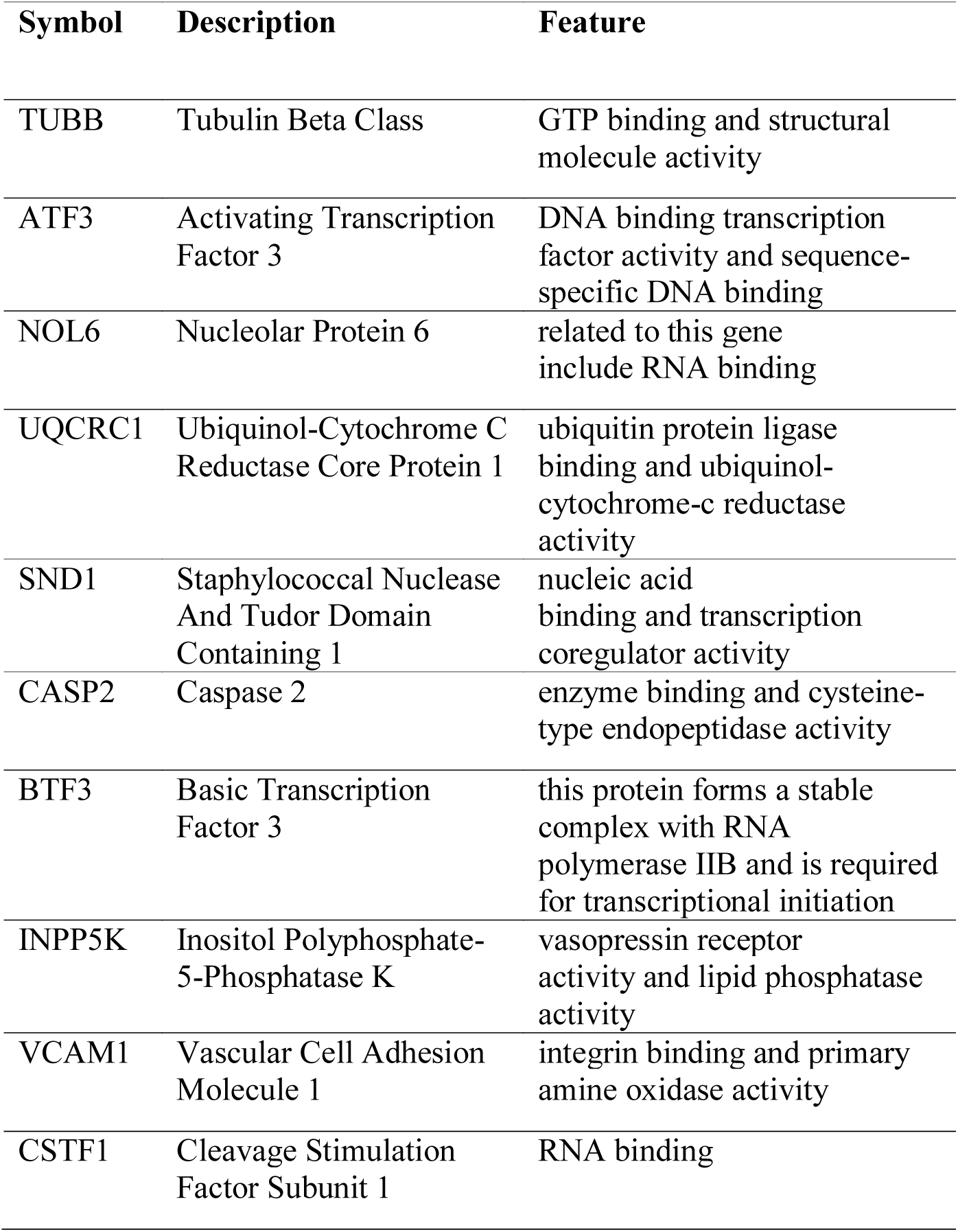
Summary of hub proteins in Alzheimer’s disease.

**Figure 3:**
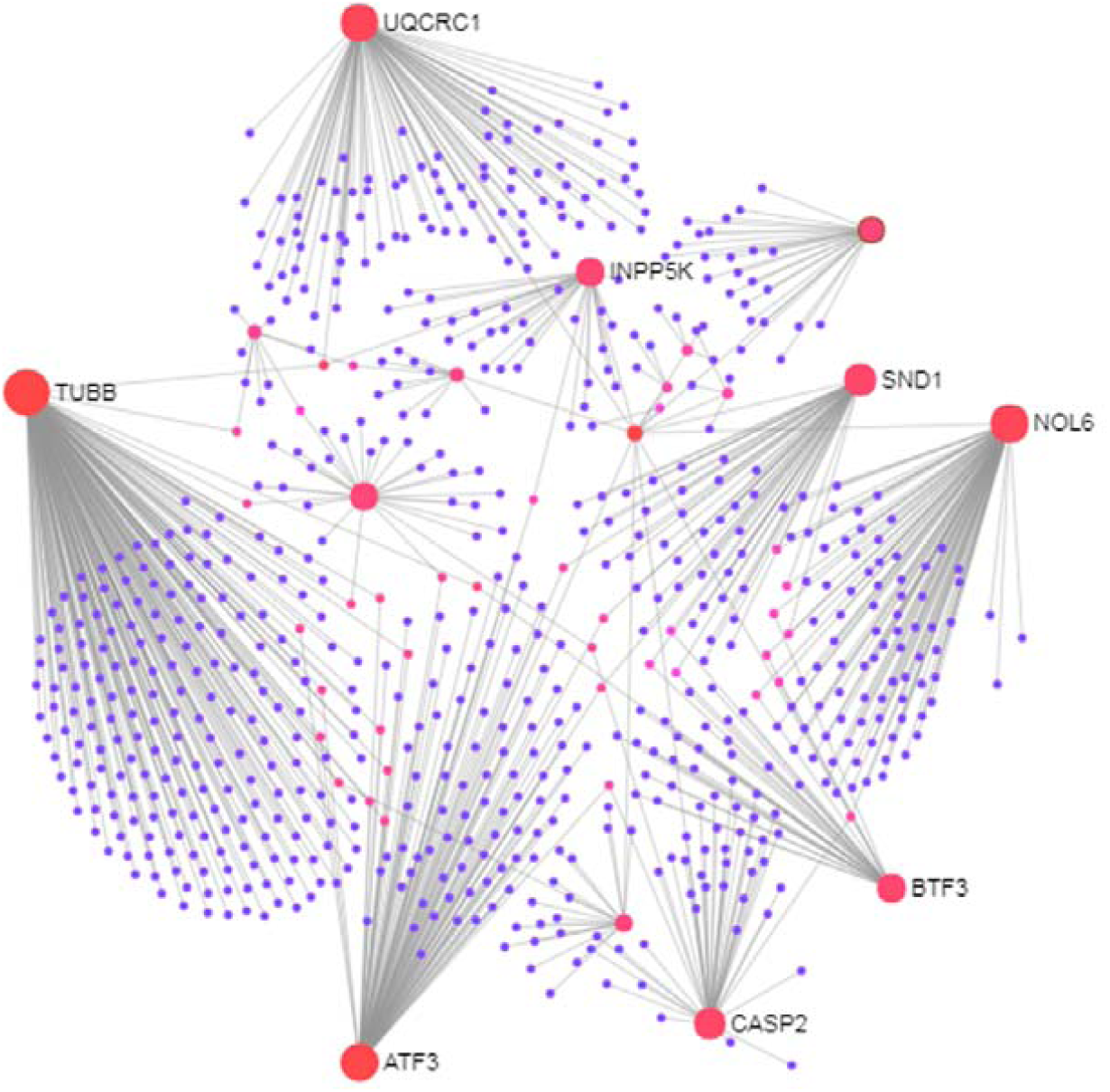
Protein-protein interaction network of the differentially expressed genes (DEGs) in Alzheimer’s disease. The nodes indicate the DEGs and the edges indicate the interactions between two genes.

### 3.4 The Regulatory Signatures of AD

We studied DEG-TF interactions and DEGs-miRNA interactions (Figure 4 and Figure 5) and detected central regulatory biomolecules (TFs and miRNAs) using topological parameters. Five TFs (FOXC1, ZNF3, GEMIN7, SMG9, and BCAP29) and 10 miRNAs (mir-20a-5p, mir-93-5p, mir-16-5p, let-7b-5p, mir-708-5p, mir-24-3p, mir-26b-5p, mir-17-5p, mir-4270, and mir-4441) were detected from the DEG-TF (Figure 4) and DEG-miRNA interaction (Figure 5) networks respectively (Table 4). These biomolecules regulate genes at transcriptional and post-transcriptional levels.

**Table 4:** Summary of regulatory biomolecules (TFs, miRNAs) in Alzheimer’s disease

| Symbol | Description | Feature |
| --- | --- | --- |
| FOXC1 | Forkhead Box C1 | DNA binding transcription factor activity and transcription factor binding |
| ZNF3 | Zinc Finger Protein 3 | nucleic acid binding and identical protein binding |
| GEMIN7 | Gem Nuclear Organelle Associated Protein 7 | Transport of the SLBP independent Mature mRNA and RNA transport. |
| SMG9 | SMG9, Nonsense Mediated MRNA Decay Factor | Viral mRNA Translation and Gene Expression. |
| BCAP29 | B Cell Receptor Associated Protein 29 | B Cell Receptor Signaling Pathway (sino) and AKT Signaling Pathway |
| miRNAs |  |  |
| mir-20a-5p | MicroRNA 20 | Parkinson's Disease Pathway and cancer |
| mir-93-5p | MicroRNA 93 | miRNAs involved in DNA damage response |
| mir-16-5p | MicroRNA 16 | sudden infant death syndrome susceptibility pathways and in cancer |
| let-7b-5p | MicroRNA let-7b | MicroRNAs in cancer and metastatic brain tumor |
| mir-708-5p | MicroRNA 708 | respiratory electron transport, ATP synthesis by chemiosmotic coupling, and heat production by uncoupling proteins. |
| mir-24-3p | MicroRNA 24 | Peptide hormone metabolism and hepatitis C and hepatocellular carcinoma |
| mir-26b-5p | MicroRNA 26 | Parkinson's Disease Pathway and cancer |
| mir-17-5p | MicroRNA 17 | Parkinson's Disease Pathway and cancer |
| mir-4270 | MicroRNA 4270 | affiliated with the miRNA class |
| mir-4441 | MicroRNA 4441 | affiliated with the miRNA class |

**Figure 4:**
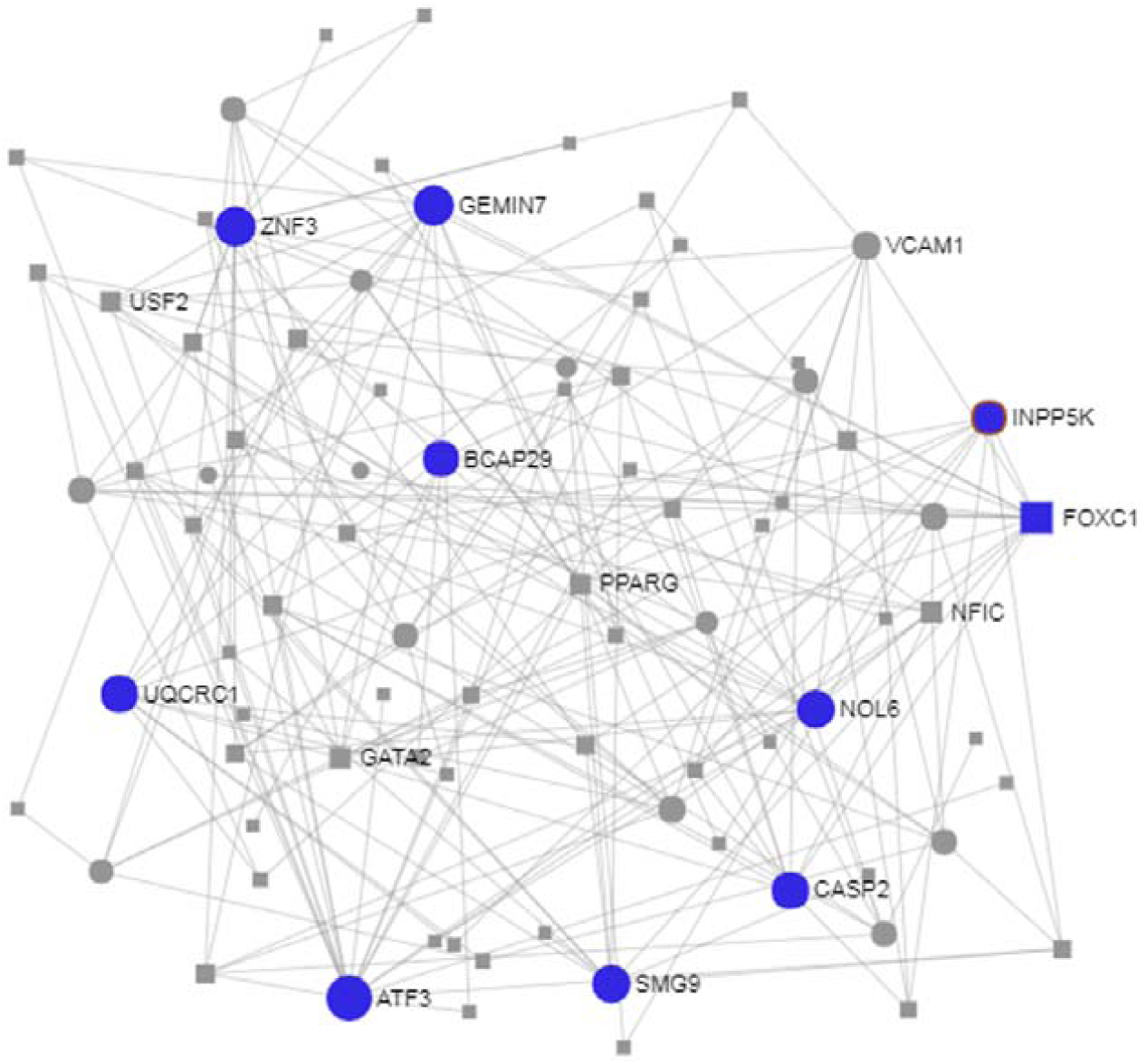
Construction of regulatory networks of DEG-TF interactions.

**Figure 5:**
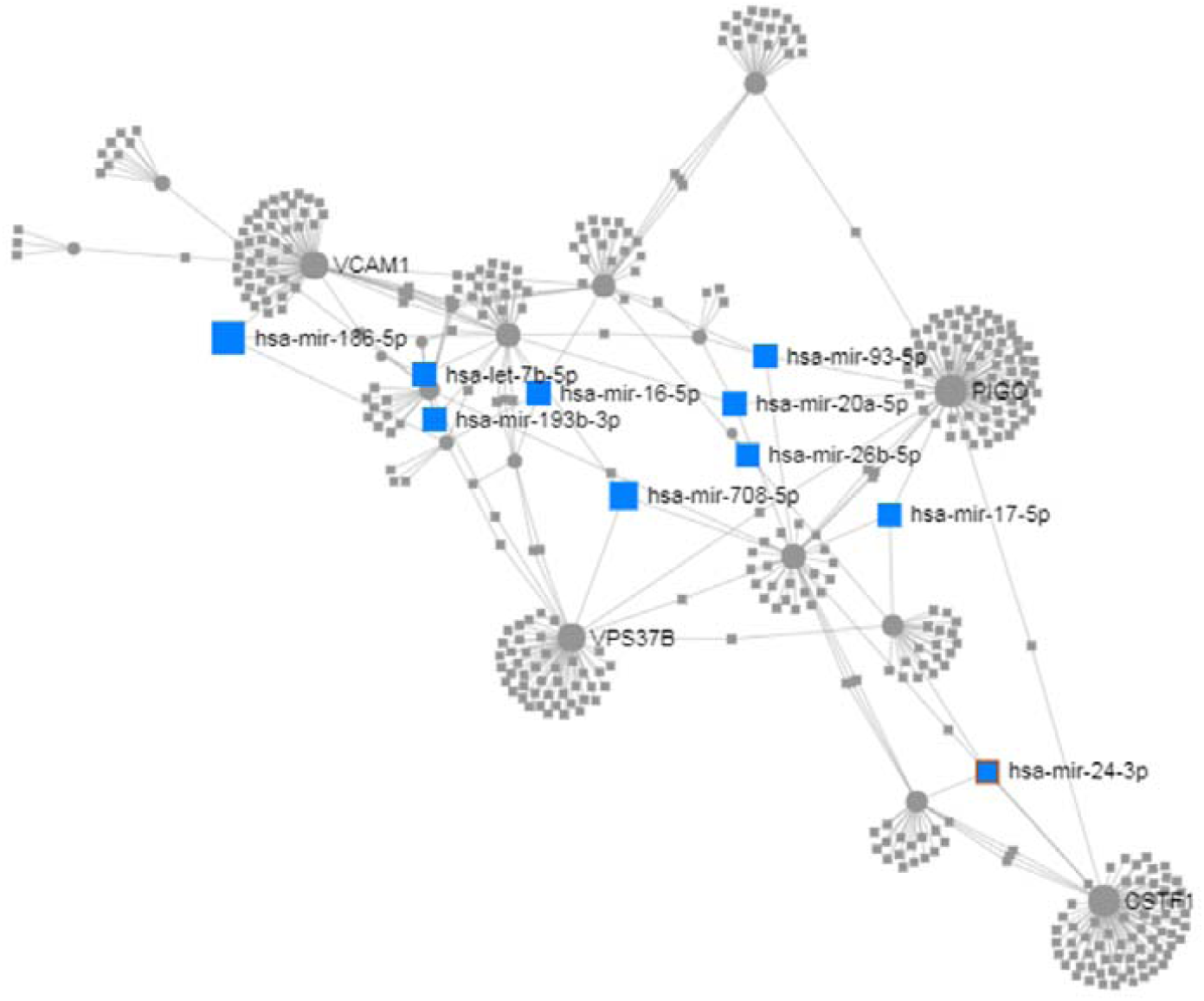
Construction of regulatory networks of DEG-miRNA interactions.

### 3.5 Epigenomic Signatures of AD: Histone Modification Sites

To provide insights regarding the epigenetic regulation of genes of interest, we studied possible histone modifications patterns. We observed that eight hub genes and TFs possessed one or more histone modification sites (Table 5).

**Table 5:** Histone modification patterns (obtained from HHMD) of novel hub genes and TFs with respect to the already known histone modification sites in neurodegenerative diseases.

| Official symbol of Biomolecules | RefSeq ID | Histone modification sites already known in neurodegenerative diseases |  |  |  |  |
| --- | --- | --- | --- | --- | --- | --- |
|  |  | H3K27 | H3K4 | H3K9 | H3K9/H4K20 | H4R3 |
| *Hub protein* |  |  |  |  |  |  |
| TUBB | NM_178014 | ✓ | ✓ | ✓ | ✓ | ✓ |
| ATF3 | NM_001030287 | ✓ | ✓ | ✓ | ✓ | ✓ |
| NOL6 | NM_139235 | ✓ | ✓ | ✓ | ✓ | ✓ |
| UQCRC1 | NM_003365 | ✓ | ✓ | ✓ | ✓ | ✓ |
| SND1 | NM_014390 | ✓ | ✓ | ✓ | ✓ | ✓ |
| CASP2 | NM_032983 | ✓ | ✓ | ✓ | ✓ | ✓ |
| BTF3 | NM_001037637 | ✓ | ✓ | ✓ | ✓ | ✓ |
| INPP5K | NM_016532 | ✓ | ✓ | ✓ | ✓ | ✓ |
| VCAM1 | NM_080682 | ✓ | ✓ | ✓ | ✓ | ✓ |
| CSTF1 | NM_001324 | ✓ | ✓ | ✓ | ✓ | ✓ |
| *TFs* |  |  |  |  |  |  |
| FOXC1 | NM_001453 | ✓ | ✓ | ✓ | ✓ | ✓ |
| ZNF3 | NM_017715 | ✓ | ✓ | ✓ | ✓ | ✓ |
| GEMIN7 | NM_001007269 | ✓ | ✓ | ✓ | ✓ | ✓ |
| BCAP29 | NM_001008407 | ✓ | ✓ | ✓ | ✓ | ✓ |

### 3.6 Cross-validation of Putative Biomarkers

A cross-check of the biomarkers identified in this study with the AlzGene database revealed no overlapped of the 25 DEGs with the AD-related biomarkers in AlzGene database. The present study did, however, identify miR-26a-5p, a miRNA that has previously been identified in a study of 12 blood-cell based miRNAs signatures of AD.

### 3.7 Protein-drug Interactions

In the protein-drug interaction network, TUBB protein was associated with the drugs Vinorelbine, Vincristine, Vinblastine, Podofilox, Colchicine, Epothilone D, Epothilone B, Cyt997, Ca4p, and Zen-012 (Figure 6). These identified drugs were categorized into different classes based on therapeutic classes (Figure 7A) and further classified according to their progress in the therapeutic pipeline (Figure 7B). Employing statistical significance thresholds of the drug-protein interactions and the possible role of the targeted protein in AD pathogenesis, a set of protein-drug interactions was selected and a series of molecular docking simulations were performed to analyze the binding affinities of identified drugs with their target protein (Table 6). The resultant energetic state and docking scores indicated the presence of thermodynamically feasible confirmations for all these interactions.

**Figure 6:**
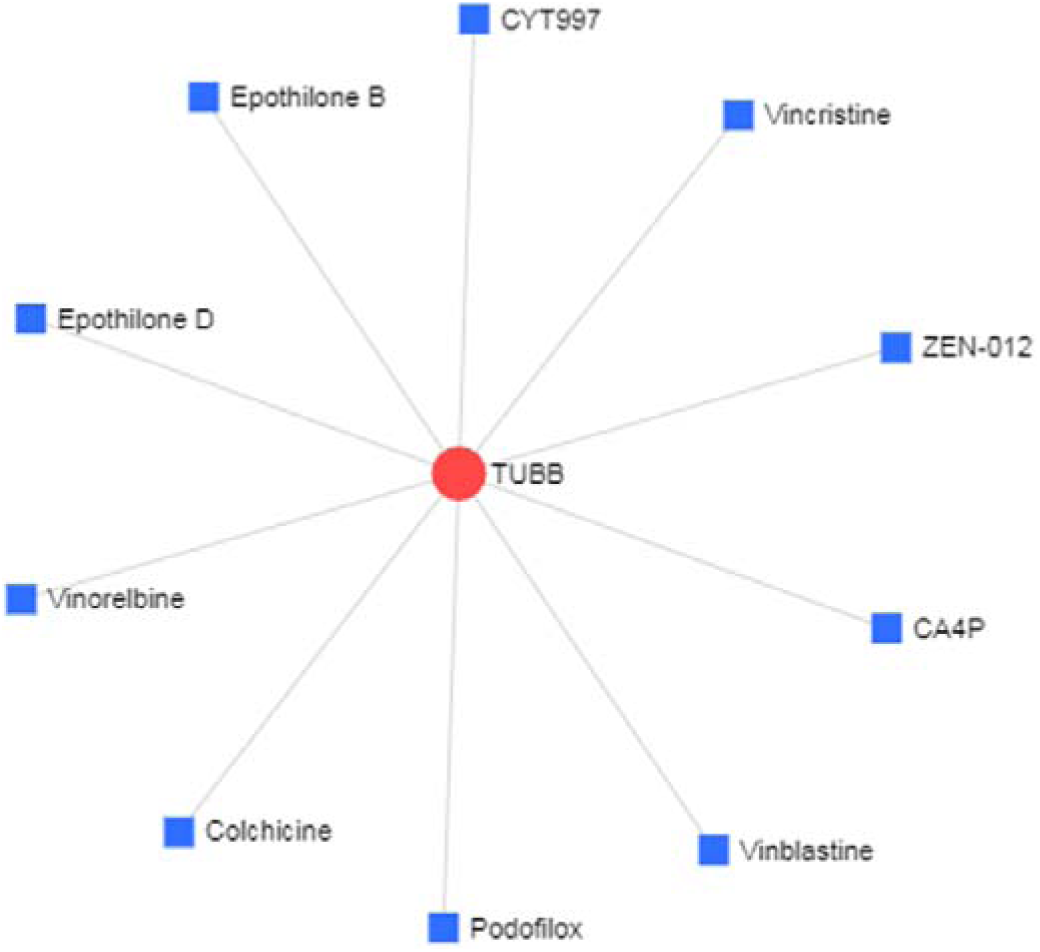
Protein-drug interaction network. The interactions between drugs and hub nodes (TUBB) are represented. The area of the node represents the degree of interaction in the network.

**Figure 7:**
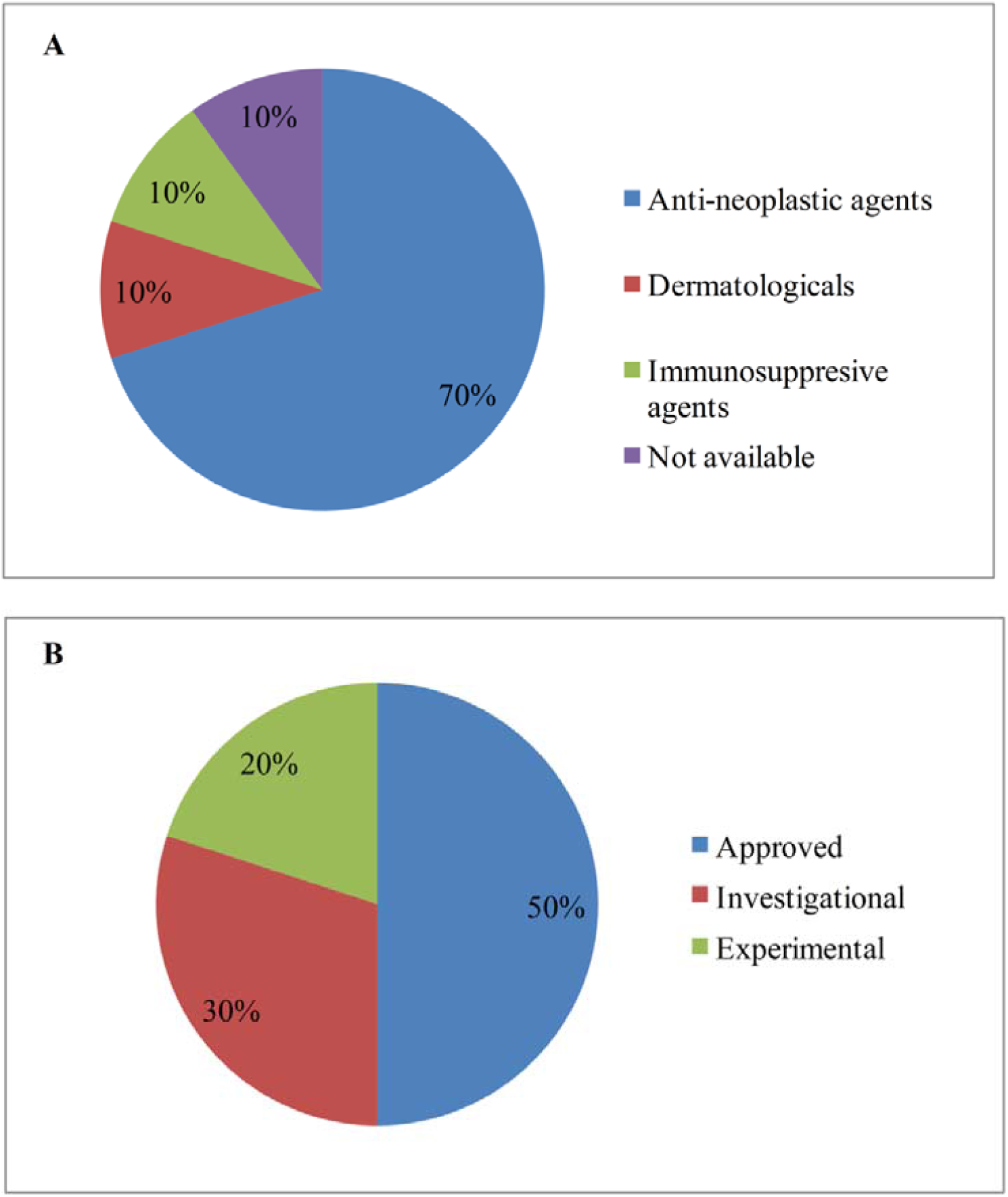
(A) Distribution of drugs into therapeutic chemical classes. (B) Classification of identified drugs according to the therapeutic pipeline.

**Table 6:**
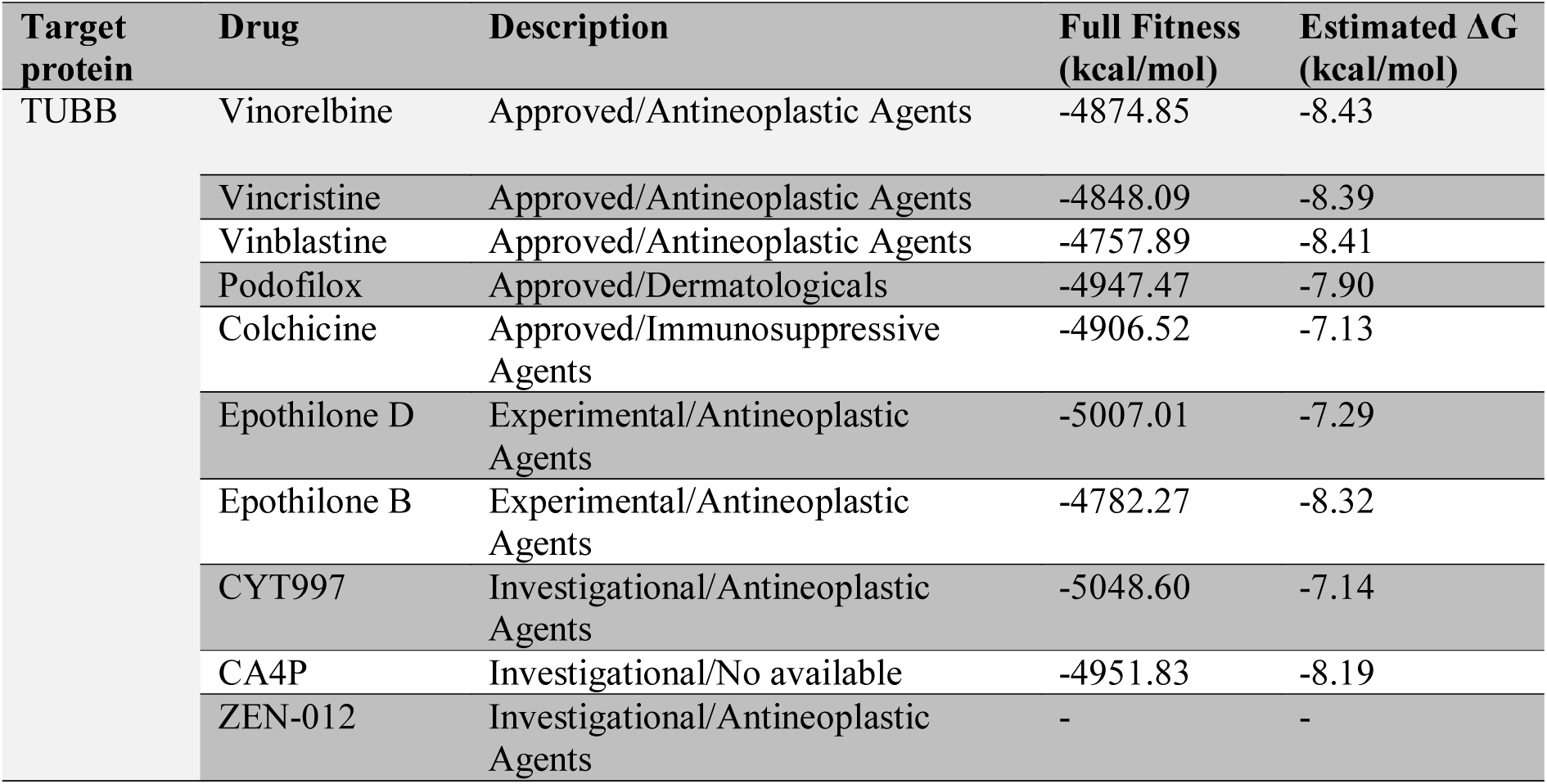
Protein-drug interactions and their binding affinity by molecular docking statistics.

### 3.8 Protein Subcellular Localization

Employing the WoLF PSORT software package we predicted the subcellular localization of proteins encoded by the DEGs in AD. The proportions of identified proteins in different subcellular compartments are displayed in Figure 8. Notably, nodes TUBB, SND1 CASP2 showed cytoplasm localization; ATF3, BTF3 were present in the nucleus; NOL6, UQCRC1, INPP5K were localized in mitochondria; VCAM1 was localized in the plasma membrane; and CSTF1 is found in both cytoplasm and nucleus.

**Figure 8:**
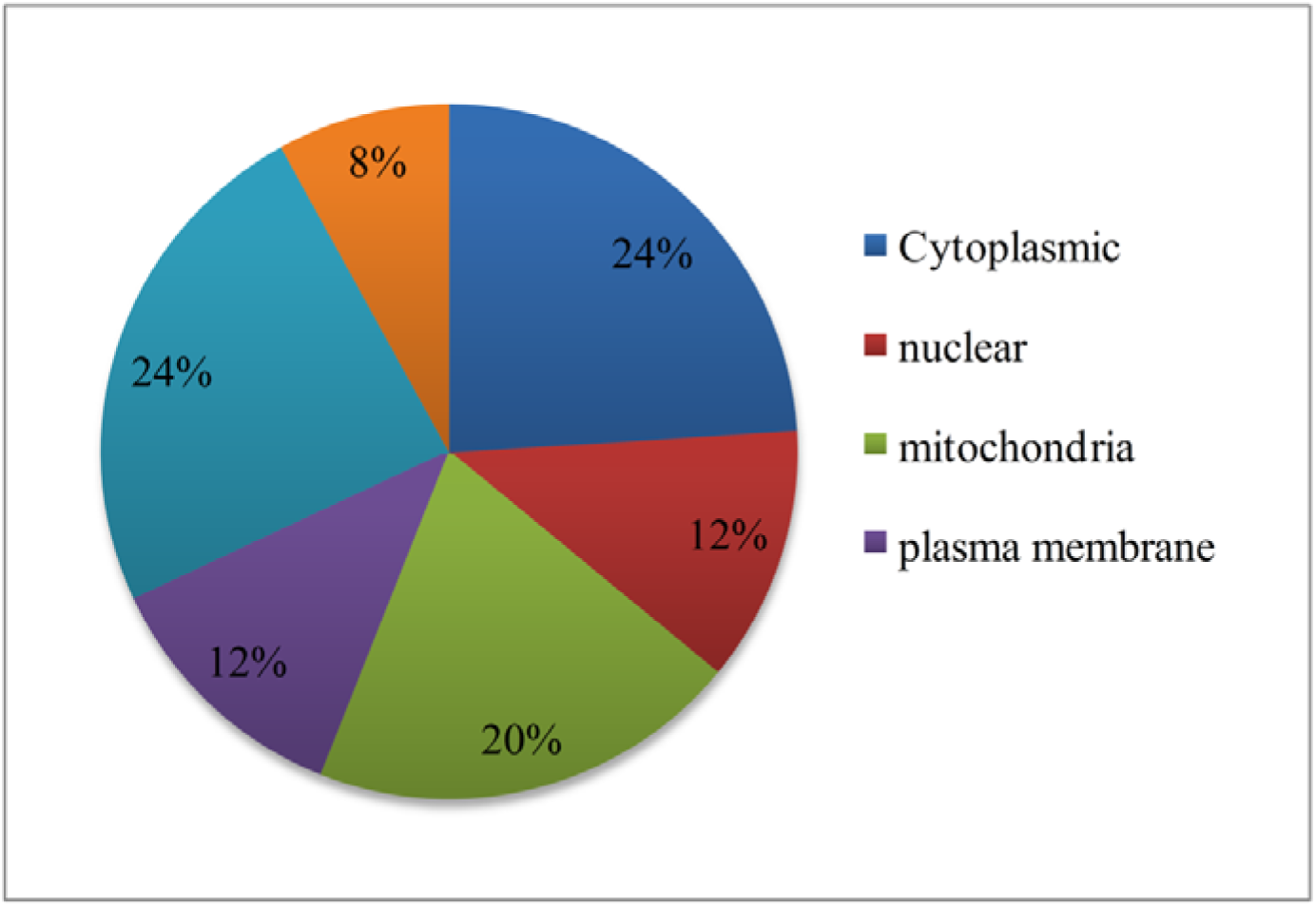
The distribution and percentages of the DEGs is shown to be the protein subcellular localization.

## 4. Discussion

The diagnosis of AD is currently performed based on neuropsychological evaluation and neuroimaging, but robust and specific biomarkers for diagnosis and prognosis of AD is an unmet challenge [11]. In this study, using a systems biology approach, we comprehensively investigated gene expression patterns evident in peripheral blood cells of AD patients and identified robust candidate biomarkers that may serve as potential biomarkers of AD. Our results may therefore provide significant insight into the process leading to AD.

Microarray data are extensively used in biomedical research and have become a major resource for identifying biomarker candidates [41]. Microarray gene expression profiling is widely used to identify DEGs in various diseases (Gov et al., 2017; Guo et al., 2017; Islam et al., 2018; Rahman et al., 2018). Analyzing gene expression patterns in the blood of AD patients revealed significant alterations in the expression profiles of 25 genes in two transcriptomic datasets. The over-representation analysis revealed AD and neurodegeneration associated molecular pathways (Oxidative phosphorylation and AD pathway) [45].

Integrative analysis of protein-protein interaction network can identify proteins that are key players in the mechanisms underlying disease [46]. Our study of the PPI network based on DEGs thus identified a number of key hub proteins (Table 2) which may indicate the onset and progression of neurodegenerative diseases. These hub proteins are associated with a number of different types of biological and pathological processes. According to the GeneCards database (https://www.genecards.org/), diseases associated with TUBB include cortical dysplasia with other brain malformations. ATF3-related pathway has been implicated in brain vascular damage [47] as well as amyloid deposition in AD [48], consistent with our observations. NOL6 has been associated with ribosome biogenesis which affects the behavior of many cells types but has not previously been linked to AD (GeneCards database). Similarly, pathways involving UQCRC1 have been implicated include cell metabolism (including respiratory electron transport), but involvement in AD has yet to be reported. The multifunctional protein SND1 is deregulated in various cancers [49] and dysregulation of CASP2, a cysteine-aspartic acid protease (caspase) family member may play a role in AD. BTF3 is a blood biomarker of AD [50]. INPP5K is associated with the disease muscular dystrophy, and intellectual disability, but no association with AD has been reported to date. Hub protein CSTF1 is involved in RNA poly-adenylation but its function is otherwise largely unknown. VCAM1 is a cell adhesion molecule important both in leukocyte and neuronal function and its expression is associated with AD dementia [51]. In summary, these hub genes have disease associations (seen in dbGaP and OMIM data) ich implies that they are involved in a range of pathogenic influences, but nevertheless most are poorly understood.

We also identified a number of TFs and miRNAs with altered levels of expression in AD patient blood cells. Differential expression of these molecules may provide significant information on dysregulation of gene expression in AD. Our previous study using microarray data obtained from brain also revealed FOXC1 as a dysregulated TF in AD (Rahman et al., 2018). A number of ZNF TFs have been implicated in the pathogenesis of neuronal diseases [52], but the role of ZNF3 in neurological disorders or AD is unknown. GEMN6 has been reported to be associated with AD [53], but this is poorly understood SMG9 is involved in nonsense-mediated mRNA decay but is not reported to have an AD association [54]. BCAP29 has been suggested as diagnostic biomarkers in peripheral blood gene expression analysis [55].

miRNAs are single stranded non-coding RNAs that can silence gene activities by targeting their transcripts [19] [56]. Dysregulated miRNAs have been reported in the development of AD [19]. Since miRNAs may serve as biomarkers for diagnosis and therapeutic target for breakthrough treatment strategies in the AD [57]; therefore, we identified the miRNAs as regulatory component of the DEGs. mir-20a-5p identified in our study is associated with aging [58] and has also been proposed as peripheral blood biomarker in AD [10]. The upregulation of mir-93-5p has been reported as biomarkers in the serum of AD patients [10]. The mir-16-5p and mir-708-5p have also been identified as differentially expressed in cerebrospinal fluid cells in AD [59,60]. The let-7b-5p has previously been down-regulated in AD as a member of 12 blood based miRNA biomarker signature [61]. The dysregulation of mir-24-3p was also reported in lung cancer [10]. Decreased expression of mir-24-3p was identified as candidate that can discriminate AD from control CSF [62]. Mir-26b-5p is also down-regulated in AD as a member of 12 blood based miRNA biomarker signature [50]. Elevated levels of miR-26-b may thus contribute to the AD neuronal pathology, and this may be reflected also in blood cell functions dependent upon mir-26b-5p [63]. Mir-17-5p was found to be deregulated in circulatory biofluids in lung cancer [10] and was dysregulated in the AD [64]. In contrast, a role for mir-4270 and mir-4441 in the AD has not been reported previously.

To study the epigenetic regulation of the hub genes, we investigated histone modification sites found in the coding genes of biomolecules implicated with neurodegenerative disease [65], identifying a range of sites, summarized in Table 4. While these are also poorly understood it raises the possibility that their alteration may be a major means of regulation for these genes that warrant further investigation, and that these genes are under tight control by a number of regulatory processes. Given the importance of these hub genes, and the possibility that they may exert sufficient influence in pathogenic processes in AD (and AD-affected blood cells) we studied the protein-drug interactions to identify drugs that may influence them. A total 10 drugs were identified from the interaction network. A number of molecular-modelling based methods are used in pharmaceutical research to investigate complex biological systems; in particular, molecular docking methods broadly used in rational drug design to elucidate the ligand conformation within binding sites of the target protein. These molecular docking methods estimate free energy by evaluating critical phenomena involved in the intermolecular recognition process in ligand-receptor binding [66]. Consequently, in the present study, we evaluated the binding mode of the ligands/drugs with the target protein TUBB and energetically stable conformations obtained from existing databases. In this way we discovered associations between the identified drugs and our putative AD markers that indicate that they may influence important pathways influencing disease progression, however it should be noted that the consequences of biomarker blockade is unclear from our work are require further investigations. We would also emphasize that our biomarkers are blood cell based and as yet we do not know how these related to brain biology. Nevertheless, many of these drugs have been evaluated for effects on tissues as part of their clinical evaluation and it may be useful to evaluate what is known of their effects on neural cells.

## 5. Conclusion

In the present study, we have analyzed the blood-based transcriptomics profiles using integrative multi-omics analyses to reveal systems level biomarkers at the protein (hub proteins, TFs) and RNA level (mRNAs, miRNAs). A number of key hub genes were significantly enriched in pathways involved in leukocyte transendothelial migration, oxidative phosphorylation, Parkinson’s disease, cell adhesion molecules, non-alcoholic fatty liver disease, AD, and Huntington’s disease. We also identified significant hub proteins TFs and miRNAs that regulate the function of the DEGs in the AD. These biomolecules may be considered candidate systems biomarkers at the protein level and RNA levels. The hub genes possess several histone modification sites of interest. Thus, we have identified potential peripheral biomarkers for AD which can be detected as transcripts in blood cells, and these warrant clinical investigations in AD patients to evaluate their utility. The nature of these biomarkers and the pathways they participate in may reveal new aspects of AD development and progression since these biomarkers are evident in cells outside the central nervous system.

## Acknowledgment

We are grateful to our colleagues for their comments and suggestion about the manuscript.

